# Asymmetric second-generation genomic incompatibility in interspecific crosses between *Ciona robusta* and *Ciona intestinalis*

**DOI:** 10.1101/2020.03.04.976837

**Authors:** Naoyuki Ohta, Nicole Kaplan, James Tyler Ng, Basile Jules Gravez, Lionel Christiaen

## Abstract

Reproductive isolation is central to speciation, but interspecific crosses between two closely related species can produce viable and fertile hybrids. Two different species in the tunicate genus *Ciona*, *Ciona robusta* and *Ciona intestinalis* can produce hybrids. However, wild sympatric populations display limited gene flow, suggesting the existence of obstacles to interspecific reproduction that remain unknown. Here, we took advantage of a closed inland culture system to cross *C. robusta* with *C. intestinalis* and established F1 and F2 hybrids. We monitored post-embryonic development, survival, and sexual maturation to further probe the physiological mechanisms underlying reproductive isolation. Partial viability of first and second generation hybrids indicated that both pre- and postzygotic mechanisms contributed to genomic incompatibilities in hybrids. Asymmetrical second generation inviability and infertility suggested that interspecific genomic incompatibilities involved interactions between the maternal, zygotic and mitochondrial genomes during development. This study paves the way to quantitative genetic approaches to study the mechanisms underlying genomic incompatibilities and other complex traits in the genome-enabled *Ciona* model.

## Introduction

Reproductive isolation is central to speciation (Mayr 1963; De Queiroz 2007), and results from the emergence of intrinsic or extrinsic barriers to reproduction that limit gene flow between populations (Seehausen *et al.* 2014). Prezygotic mechanisms of reproductive isolation, including habitat segregation, phenological and sexual isolation, hinder mating and/or fertilization between individuals of different species or populations undergoing speciation. Even when individuals overcome prezygotic obstacles to reproduction, postzygotic mechanisms can prevent growth and sexual maturation of hybrids. Intrinsic mechanisms of postzygotic reproductive isolation are referred to as genomic incompatibilities, also known as Bateson-Dobzhansky-Muller incompatibility (BDMI; (Orr 1996; Orr and Presgraves 2000; Presgraves 2010; Cutter 2012)). In addition, extrinsic postzygotic reproductive barriers, including interactions with the environment and with other individuals, can reduce the viability and/or fertility of the hybrid offspring. Despite obstacles to reproduction, individuals from closely related species occasionally produce viable and fertile hybrids, which in turn impact gene flow and species evolution, particularly through introgressions at specific loci between distinct genomes (Roux *et al.* 2013; Bouchemousse *et al.* 2016b).

Within the ascidian genus *Ciona*, distinct type A and type B were identified within the species *Ciona intestinalis*. These types were first thought to represent cryptic sub-species (Suzuki *et al.* 2005; Kano *et al.* 2006; Caputi *et al.* 2007; Nydam and Harrison 2010; Sato *et al.* 2012, 2014). They were more recently recognized as two distinct species, *Ciona robusta* and *Ciona intestinalis*, respectively (Brunetti *et al.* 2015). Based on molecular clock estimates, the speciation event that segregated *C. robusta* and *C. intestinalis* is thought to have occurred approximately 4 million years ago (Mya; (Nydam and Harrison 2007; Roux *et al.* 2013), following geographical separation between the Pacific and Atlantic oceans (Caputi *et al.* 2007; Bouchemousse *et al.* 2016b). However, the two species came in contact secondarily, and co-exist in the English Channel, where *C. intestinalis* is the original endemic species, while *C. robusta* is thought to have invaded the area, in through human transportation (Zhan *et al.* 2010; Nydam and Harrison 2011; Sato *et al.* 2012; Roux *et al.* 2013; Bouchemousse *et al.* 2016a). In the contact area, natural hybrids of *C. robusta* and *C. intestinalis* were found, but at very low frequencies. Furthermore, the two species displayed limited exchange of alleles (Nydam and Harrison 2011; Sato *et al.* 2012; Bouchemousse *et al.* 2016c), suggesting that mechanisms ensuring reproductive isolation largely restrict the expansion of hybrids, as well as gene flow between the two species in the contact region.

Mechanisms ensuring species-specific fertilization are important for prezygotic reproductive isolation (Mayr 1963; Seehausen *et al.* 2014; Herberg *et al.* 2018), but successful fertilization between *C. robusta* and *C. intestinalis* can routinely be obtained in the laboratory, despite indications that *C. intestinalis* sperm occasionally fails to fertilize *C. robusta* eggs (Suzuki *et al.* 2005; Sato *et al.* 2014; Bouchemousse *et al.* 2016a; Malfant *et al.* 2017). Notably, *Ciona* adults are self-incompatible hermaphrodites (Harada *et al.* 2008; Sawada *et al.* 2020), which spawn their gametes in the open water at dawn. Intrinsic prezygotic isolation would thus involve gamete recognition and/or fertilization success rather than, for example, mating behavior. Nonetheless, prezygotic reproductive isolation in *Ciona* may not suffice to explain the quasi-absence of natural hybrids and limited gene flow in the wild. Instead, it is thought that postzygotic mechanisms ensure reproductive isolation, including genomic incompatibility in the second generation hybrids. For instance, Sato and colleagues crossed F1 hybrids, produced by forcibly crossing *C. robusta* and *C. intestinalis*, and obtained backcrossed BC1 larvae (Sato *et al.* 2014). However, to our knowledge, the viability and fertility of F2 hybrids, which could provide clues about the physiological origins of the reproductive isolation between *Ciona robusta* and *Ciona intestinalis*, has not been reported.

In this study, we took advantage of a closed inland culture system to cross *C. robusta* and *C. intestinalis*, and maintain hybrids through multiple generations. We assayed survival, growth and sexual maturation, to further evaluate pre- and postzygotic mechanisms of reproductive isolation between *C. robusta* and *intestinalis*. Our observations indicate that F1 and F2 hybrids have reduced fitness compared to *C. robusta*, suggesting the existence of pre- and postzygotic mechanisms of reproductive isolation. Additionally, we report evidence of asymmetric second generation incompatibilities emerging from reciprocal crosses, as commonly observed in numerous plant and animal taxa.

## Materials and Methods

### Animals

Wild-type *Ciona robusta* (*C. intestinalis* type A) and *Ciona intestinalis* (*C. intestinalis* type B) adults were collected in San Diego (CA) and Woods Hole (MA), respectively, and are within the range of known distribution for these species (Nydam and Harrison 2007; Caputi *et al.* 2007; Bouchemousse *et al.* 2016b). We confirmed species identification using established phenotypic criteria (Sato *et al.* 2012). Sperm and eggs were surgically obtained from mature animals, and used for controlled *in vitro* fertilizations to produce F1 generation animals, using established protocols (Christiaen *et al.* 2009). We cultured all animals at 18°C (Sanyo, MIR-154), a permissive temperature for both species, as well as for F1 hybrids (Sato *et al.* 2014; Malfant *et al.* 2017). Juveniles were kept in Petri dishes at 18°C until 28 days post fertilization (dpf). We changed the buffered artificial sea water in the dishes and fed animals every other day. The young animals were transferred into a closed inland culture system at 28 dpf. We measured survival rate by counting the number of live animals in each Petri over time, and measured the size of each living individual from the tip of the siphon to the end of the body. The data was analyzed using Microsoft Office Excel and R. We dissected mature F1 animals to obtain sperm and/or eggs and generated F2 animals by controlled *in vitro* fertilization.

### Algae culture

We essentially followed an established protocol (Joly *et al.* 2007). We used two strains of microalgae, *Chaetoceros gracilis* and *Isochrysis galbana* (aka T.iso) as food for *Ciona* juveniles and adults, 10^7^ to 10^8^ cells for each Petri dish, and 10^9^ to 10^10^ cells for tanks. Stock, starter, and scale-up cultures of algae were kept in 250mL, 500mL and 2L flasks, respectively. Terminal food cultures were kept in 10L carboys. The flasks and carboys were maintained under constant light (Marineland), and were shaken once a day to prevent sedimentation. The cultures were inoculated every 10 to 14 days. Half of the cultures were diluted with autoclaved artificial sea water (Bio-Actif Salt, Tropic Marin) for the next round of cultures. We added 1mL Conway medium (Bouquet *et al.* 2009; Martí-Solans *et al.* 2015) and 1g sodium bicarbonate (Sigma Aldrich) per 1L artificial sea water, instead of bubbling CO_2_, to scale up intermediate and terminal food cultures. We added 1mL silicate solution (40g/L metasilicate sodium, Fisher Scientific) per 1L artificial sea water for scale-up and ongoing food cultures of *Chaetoceros*. Conway medium contains: 45g/L EDTA (C_10_H_16_N_2_O_8_, Acros Organics) 100g/L Sodium nitrate (NaNO_3_, Acros Organics), 33.3g/L Boric acid (H_3_BO_3_, Fisher Scientific), 20g/L Sodium dihydrogen phosphate (NaH_2_PO, Acros Organics), 1.5g/L Manganese (II) chloride (MnC_12_•4H_2_O, Acros Organics), 1.3g/L iron (III) chloride (FeCl_3_•6H_2_O, Acros Organics), 21mg/L Zinc chloride (ZnCl_2_, Acros Organics), 20mg/L Cobalt (II) chloride (CoCl_2_•6H_2_O, Acros Organics), 10mg/L Ammonium heptamolybdate ((NH_4_)_6_Mo_7_O_24_•4H_2_O, Acros Organics), 21mg/L Copper (II) sulphate (CuSO_4_•5H_2_O, Acros Organics), 10mg/L Thiamine (Acros Organics), 10mg/L Cyanocobalamin (Acros Organics), 200μg/L Biotin (Fisher Scientific).

### System maintenance

The culture system held 20L glass aquarium tanks (Carolina), 5L polypropylene beakers (Midland Scientific) and 2L polycarbonate aquarium tanks (Eisco), which each could hold 16, 4 and 2 Petri dishes, respectively (Supplemental Figure S1A-C). The 20L tanks, 5L beakers and 2L small tanks were set in an 18°C chamber. Sea water (Bio-Actif Salt, Tropic Marin) was controlled by bio-balls (Biomate, Lifegard Aquatics) and bacteria (BioDigest, Prodibio), and salinity was set at 34 ppt. The 20L tanks were cleaned twice a week, and the 5L beakers and the 2L small tanks were cleaned three times a week. The 20L tanks, 5L beakers and 2L small tanks each efficiently supported animal growth and sexual maturation (Supplemental Figure S1D-E).

### Genotyping

F1 juveniles were taken from Petri dishes and digested by proteinase (Thermo Fisher Science) to obtain genomic DNA as described previously (Ohta *et al.* 2010). Oral siphon and sperm were surgically obtained from mature animals, and processed for genomic DNA extraction by QIAamp DNA Micro Kit (Qiagen), or digested with proteinase. The genomic DNA was used for PCR amplification (35 cycles of 95°C 30’’, 58°C 30’’, 72°C 1’) of target regions with Ex Taq HS DNA polymerase (Takara Bio). The PCR products were purified enzymatically with ExoSAP-IT Express PCR product cleanup (Thermo Fisher Scientific), or by NucleoSpin Gel and PCR clean-up (Macherey-Nagel). PCR products were sequenced by Genewiz. The primers used in this study are summarized in Supplemental Table S1.

The authors affirm that all data necessary for confirming the conclusions of the article are present within the article, figures, and tables.

## Results

### Reciprocal crosses between *Ciona robusta* and *C. intestinalis* produce hybrids

In order to cross *Ciona robusta* and *Ciona intestinalis*, we obtained mature animals from San Diego (CA) and Woods Hole (MA), respectively (Figure 1A). Using six isolated batches of sperm and eggs from each species, we performed homotypic and heterotypic crosses by *in vitro* fertilization to obtain twenty-four combinations of four types of animals in three separate partial diallels: the parental strains *C. robusta* and *C. intestinalis*, and reciprocal F1 hybrids, which we termed RxI, and IxR, for hybrids obtained from *C. robusta* sperm and *C. intestinalis* eggs, or *C. intestinalis* sperm and *C. robusta* eggs, respectively (Figure 1B). We obtained hundreds of swimming larvae from each cross (Figure 2A-D), and did not estimate fertilization rates or the proportion of hatched larvae, although this contrasts with previous studies, which suggested that *C. robusta* oocytes were largely refractory to fertilization by *C. intestinalis* sperm (Suzuki *et al.* 2005; Bouchemousse *et al.* 2016a; Malfant *et al.* 2018). Further work will be required to determine whether these discrepancies stem from biological and/or experimental differences between studies.

**Figure 1.**
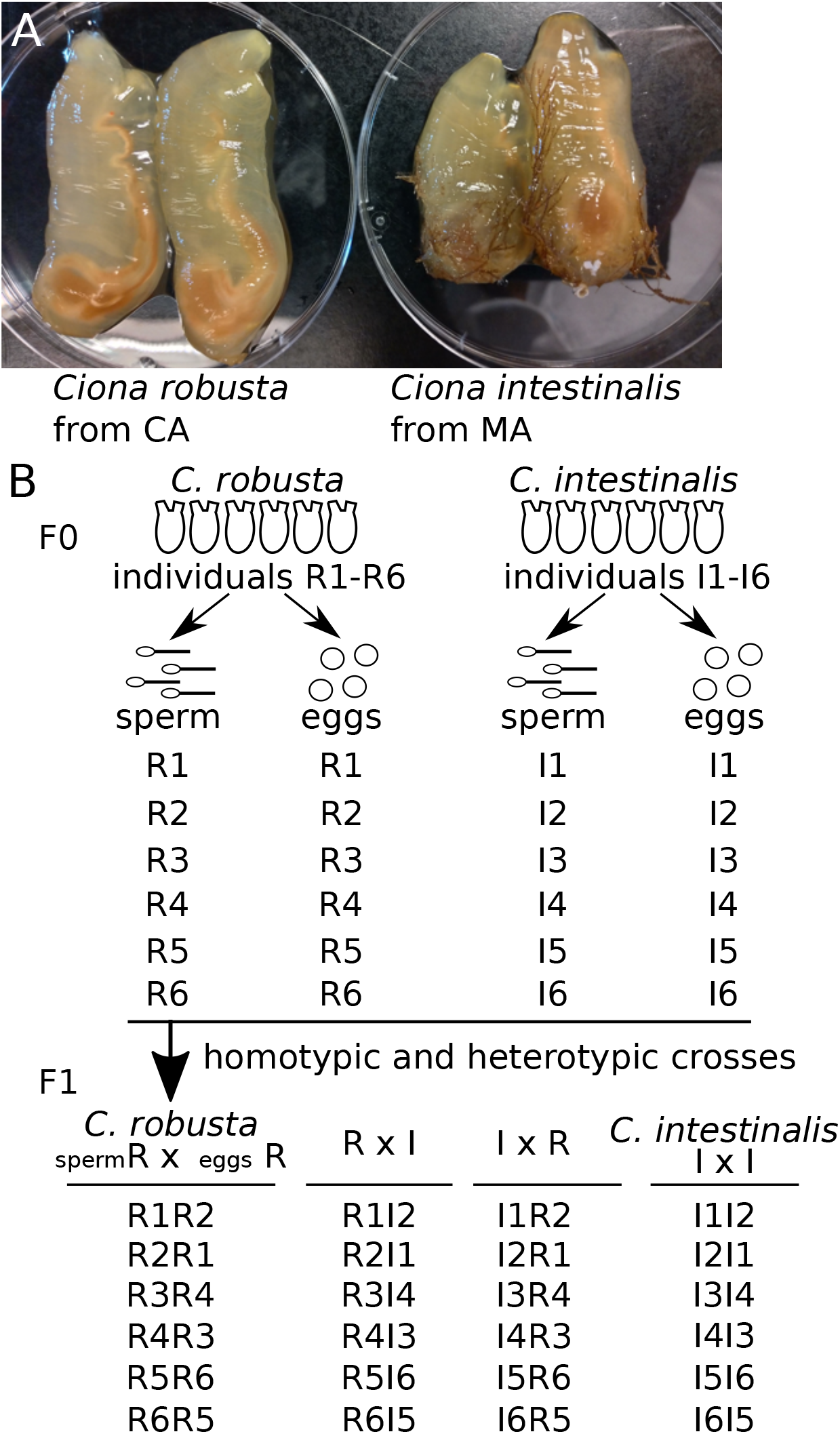
Crossing animals to make genetic hybrids. (A) Mature animals of *Ciona robusta* and *Ciona intestinalis* were collected in San Diego (CA) and Woods Hole (MA), respectively. The animals were in 100 mm Petri dishes. (B) Six animals were dissected to obtain sperm R1 to R6 and eggs R1 to R6 of *C. robusta* and sperm I1 to I6 and eggs I1 to I6 of *C. intestinalis*. These sperm and the eggs were homo- and heterotypic crossed to make *C. robusta* (sperm R x eggs R), RxI hybrid, IxR hybrid and *C. intestinalis* (IxI) types.

**Figure 2.**
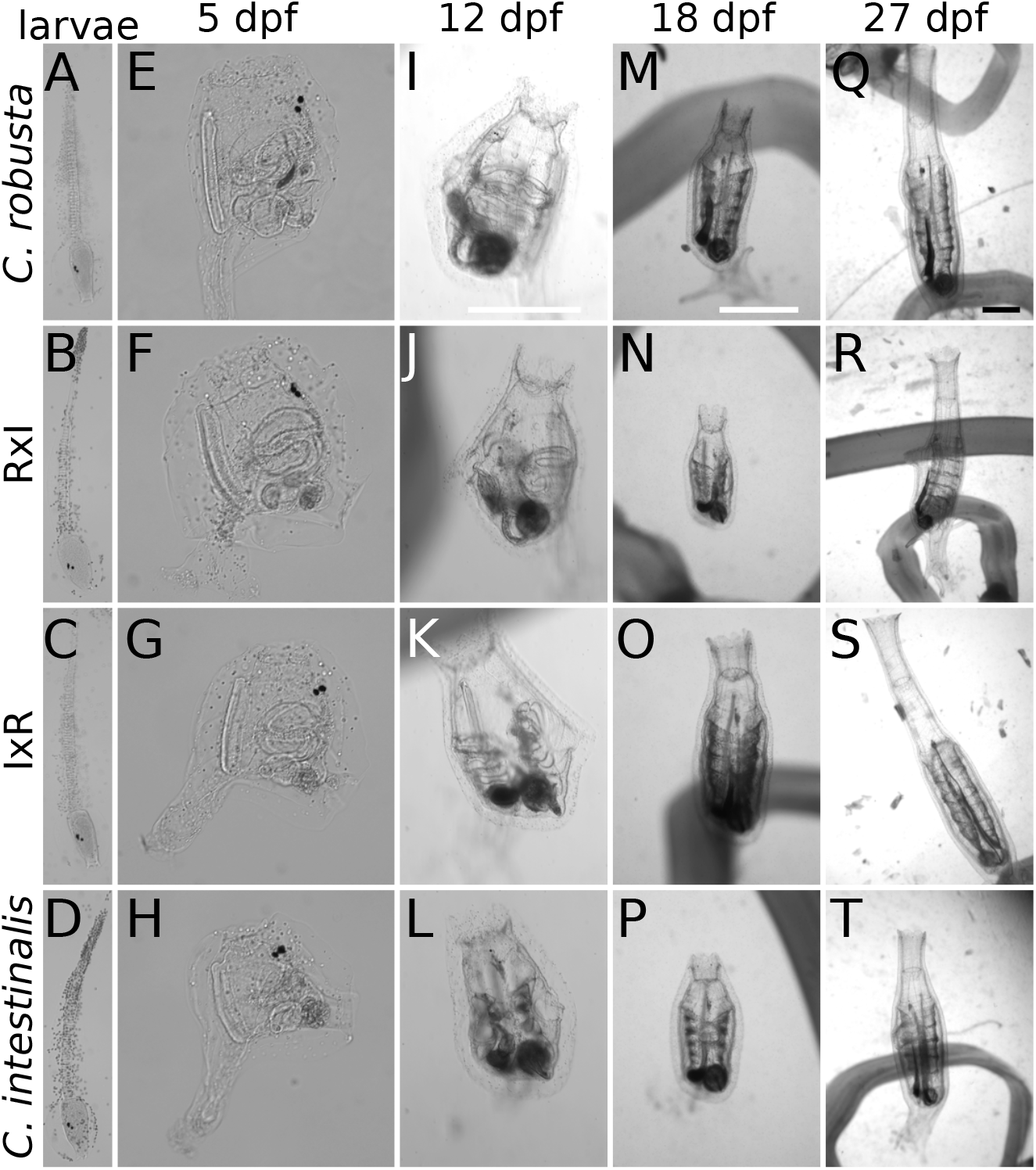
Development of F1 animals. (A-H) Images taken after fixation in formamide of: swimming larvae (A-D) and 5 dpf juveniles (E-H). (I-T) Images of living animals were taken under a microscope of: 12 dpf juveniles (I-L), 18 dpf young adults (M-P) and 27 dpf young adults (Q-T). Scale bars in I, M and Q show 1 mm.

We monitored development following hatching, settlement, metamorphosis and initial growth, and did not observe obvious differences between the four types, although this cursory analysis may have missed subtle quantitative variability (Figure 2E-T). We measured survival rates from 5 to 50 days post fertilization (dpf) by counting the number of animals in each Petri dish (Figure 3A-B). About 70% of animals in all four conditions survived to 50 dpf (Figure 3A-B), and an ANOVA did not show significant differences in survival rate between the four types at 26 and 50 dpf, except for between *C. robusta* and RxI hybrid at 50 dpf (Figure 3B). Notably, there were no significant differences in the survival rate between F1 RxI and IxR hybrids at 26 or 50 dpf. We monitored the size of animals from 18 dpf to 50 dpf, while keeping the feeding regime constant across conditions (Figure 3C-D). Here too, an ANOVA did not reveal significant differences in the size of F1 hybrids at 26 dpf, although size significantly differed between hybrids and *C. robusta* at 50 dpf (Figure 3D). Notably, an ANOVA did not show significant size differences between F1 RxI and IxR hybrids at 26 dpf, but showed it at 50dpf. A previous study reported differences in growth rate for hybrid animals of 28 dpf (Malfant *et al.* 2018). Taken together, these observations suggest that reciprocal first generation hybrids of *C. robusta* and *C. intestinalis* are generally as healthy as the parental strains, as they did not display marked differences in post-hatching survival and growth.

**Figure 3.**
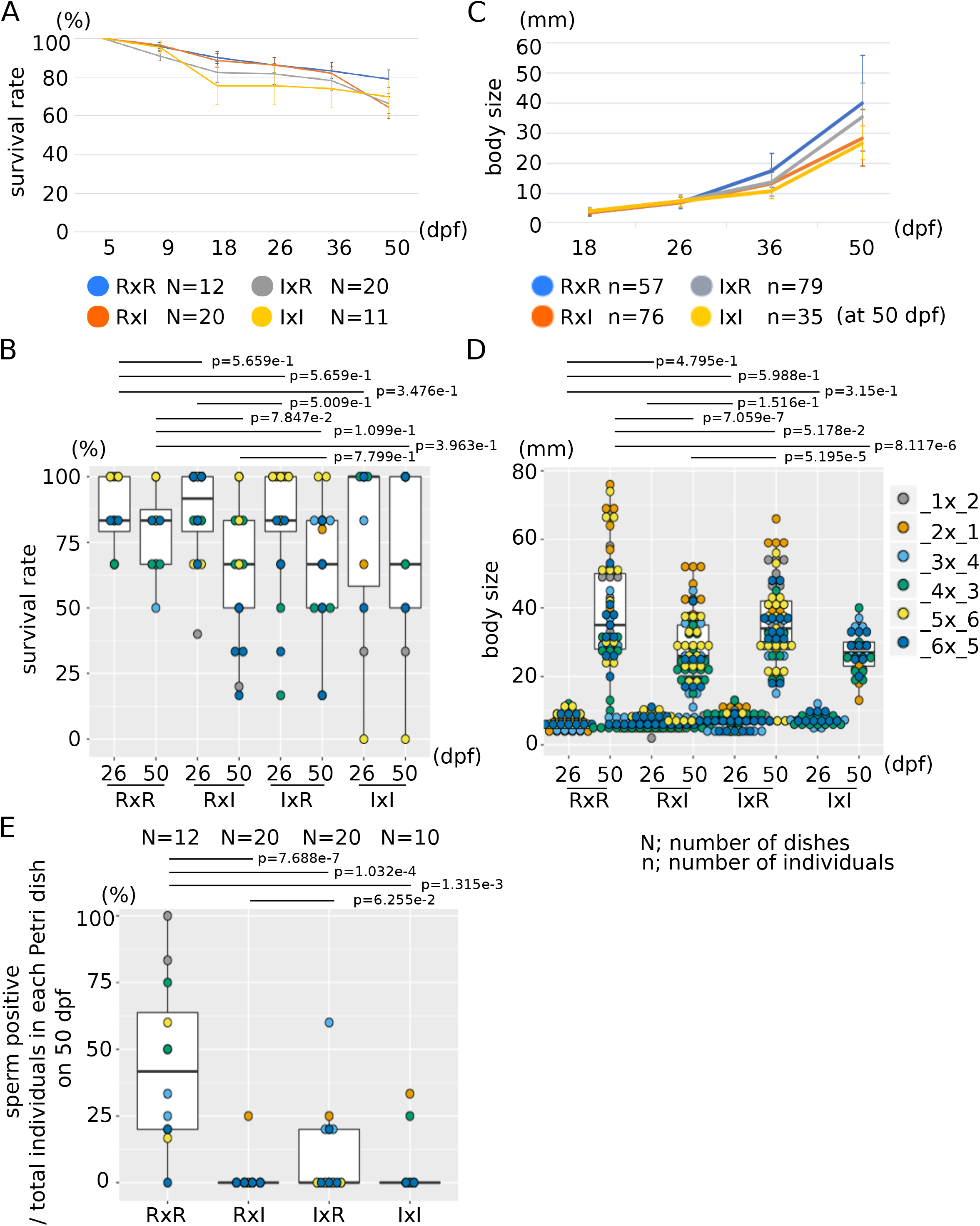
Growth of F1 animals. (A) The chart shows the average survival rate in each Petri dish. Error bars show standard deviation. (B) The dot and boxplot shows the survival rate in each Petri dish at 26 and 50 dpf. Color shows each Petri dish in each parental combination. (C) The chart shows the average size of each individual. Error bars show standard deviation. (D) The dot and boxplot shows the size of each individual at 26 and 50 dpf. Color shows each individual in each parental combination. (E) The dot and boxplot shows the ratio of animals which had sperm at 50 dpf in each Petri dish. N shows the numbers of Petri dishes. n shows the numbers of individuals at 50 dpf. p values were calculated by an ANOVA.

Next, we sought to raise *Ciona* hybrids to sexual maturity in our experimental facility. In a previous report, Sato and colleagues cultured F1 hybrids in their natural environment, the English Channel, where the two species live in sympatry, and obtained mature animals after a couple of months (Sato *et al.* 2014). Here, we took advantage of a custom inland culture system to raise and monitor animals through sexual maturation. By 50 dpf, half of the *C. robusta* individuals were producing sperm, whereas that proportion dropped significantly for the other groups of animals (Figure 3E). We kept these animals until they produced eggs and/or sperm, which we collected surgically, thus sacrificing F1 animals, to test their fertility and obtain F2 animals (Table 1).

**Table 1.** Summary F1.

|  | *C. robusta* | Rxl | lxR | *C. intestinalis* |
| --- | --- | --- | --- | --- |
| sperm+ | 14 | 16 | 21 | 7 |
| used for F2 | 6 | 13 | 18 | 6 |
| discarded | 8 | 3 | 3 | 1 |
| egg+ | 2 | 6 | 13 | 0 |
| used for F2 | 0 | 6 | 11 | 0 |
| discarded | 2 | 0 | 2 | 0 |
| dead or unknown | 7 | 39 | 33 | 0 |
| discarded | 36 | 20 | 22 | 28 |
| total | 57 | 76 | 79 | 35 |

### Phenotypes of hybrid adult animals

One obvious difference between parental species is the presence of an orange pigment organ (OPO) at the tip of the sperm duct in *C. robusta*, but not in *C. intestinali*s (Millar 1953; Hoshino and Tokioka 1967; Ohta *et al.* 2010; Sato *et al.* 2012, 2014; Tajima *et al.* 2019). F1 animals from our parental strains did recapitulate this species-specific trait (Figure 4A, A’, D and D’). For both RxI and IxR hybrids, the majority of animals had OPO at the tip of the sperm duct (Figure. 4B, B’, C, C’ and E), in agreement with a previous report (Sato *et al.* 2014), thus indicating that OPO formation is a dominant trait.

**Figure 4.**
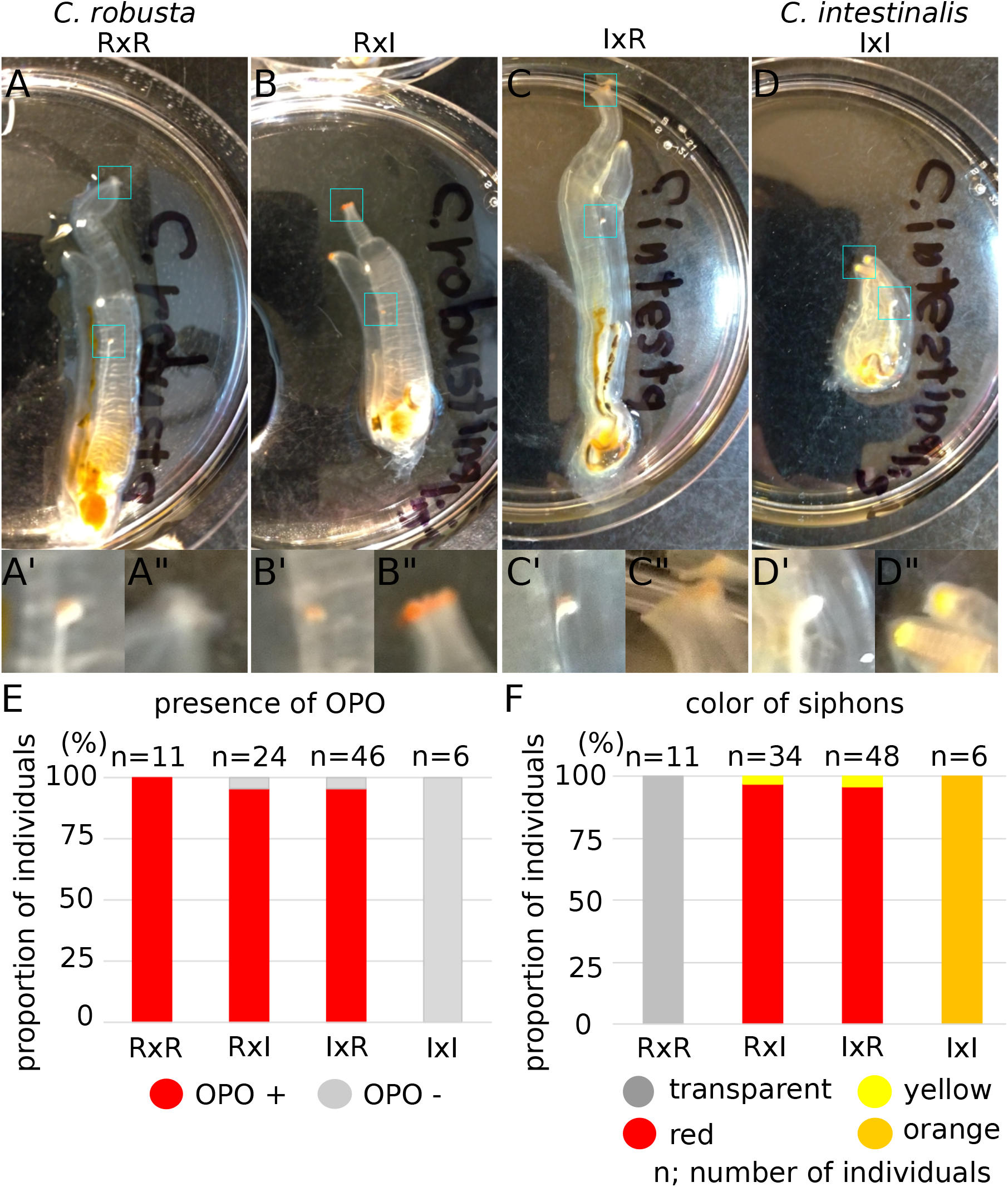
Phenotype of F1 mature animals. (A) Images were taken of F1 mature animals in 100 mm Petri dishes; *C. robusta* (A), RxI hybrid (B), IxR hybrid (C) and *C. intestinalis* (D). (A’-D’, A’’-D’’) The tip of sperm duct (A’-D’) and oral siphon (A’’-D’’) of each individual are shown in the insets. (E) The bar shows the proportion of individuals having OPO at the tip of sperm duct in F1 mature animals. (F) The bar shows the proportion of individuals having color in the rim of oral and atrial siphons in F1 mature animals. n shows the numbers of individuals.

Another character that differs between *Ciona* species is the color of siphons (Sato *et al.* 2012), whereby *C. intestinalis* has yellow and orange pigmentation around the tip of siphons that is lacking in *C. robusta* (Figure. 4A, A’’, D and D’’), although this feature was deemed quite variable and taxonomically unreliable (Brunetti *et al.* 2015). As for the OPO, the majority of RxI and IxR hybrids displayed a bright red pigmentation at the rim of oral and atrial siphons (Figure 4B, B’’, C, C’’ and F), also consistent with a previous report (Sato *et al.* 2014). The observation that siphon pigmentation displays an overdominant phenotype in hybrids is consistent with its lack of reliability for taxonomic purposes. Further work will be required to determine how proposed species-specific and taxonomically informative traits, such as tubercular prominences in the siphons (Brunetti *et al.* 2015), which we could not observe clearly, are inherited through generations of hybrids.

### Genotyping of hybrid animals

The distribution of variable traits in homo- and heterospecific crosses suggested that RxI and IxR F1 animals are *bona fide* hybrids. As a complement to phenotypic characterization, and to rule out cross-contaminations during the *in vitro* fertilization procedure, we sought to perform molecular genetics analyses to assay the distribution of species-specific marker alleles in the different strains (Suzuki *et al.* 2005; Nydam and Harrison 2007). We unsuccessfully tested two primer sets, markers 1 and 2, which were previously used to distinguish *C. robusta* and *C. intestinalis* ((Suzuki *et al.* 2005); Supplemental Figure S2A-C). However, sequence differences between the PCR products distinguished between species-specific alleles (Supplemental Figure S2D). As an alternative, we used a primer set designed at *Myosin light chain 2/5/10* (*Myl2/5/10*; KH.C8.239) locus, which could distinguish *C. robusta* and *C. intestinalis* alleles by the size difference of PCR products (Figure 5A, Supplemental Figure S3A-B). Sequencing amplicons showed conserved 6^th^ and 7^th^ exons, but an indel in the 6^th^ intron that distinguished alleles from different species (Supplemental Figure S3C). We isolated genomic DNA from three F1 juvenile individuals from each type. Six juveniles of either *C. robusta* or *C. intestinalis* yielded single bands, albeit of higher molecular weight for the latter (Supplemental Figure S3D). By contrast, six juveniles of either RxI or IxR crosses yielded double bands, showing that these animals had both *C. robusta* and *C. intestinalis Myl2/5/10* alleles, and were indeed hybrids. Consistent with electrophoresis patterns, sequence analysis revealed single alleles for either *C. robusta* or *C. intestinalis*, whereas F1 RxI and IxR hybrids produced a mixture of *C. robusta*- and *C. intestinalis*-specific sequences (Supplemental Figure S3E). Of note, genomic DNA from both somatic tissue and gametes yielded similar results, whereby homotypic *C. robusta* and *C. intestinalis* produced single PCR bands in different sizes, while those of both hybrids produced double PCR bands (Figure 5B). Taken together with the results of phenotypic observations, genotyping data indicated that F1 RxI and IxR animals were *bona fide* hybrids.

**Figure 5.**
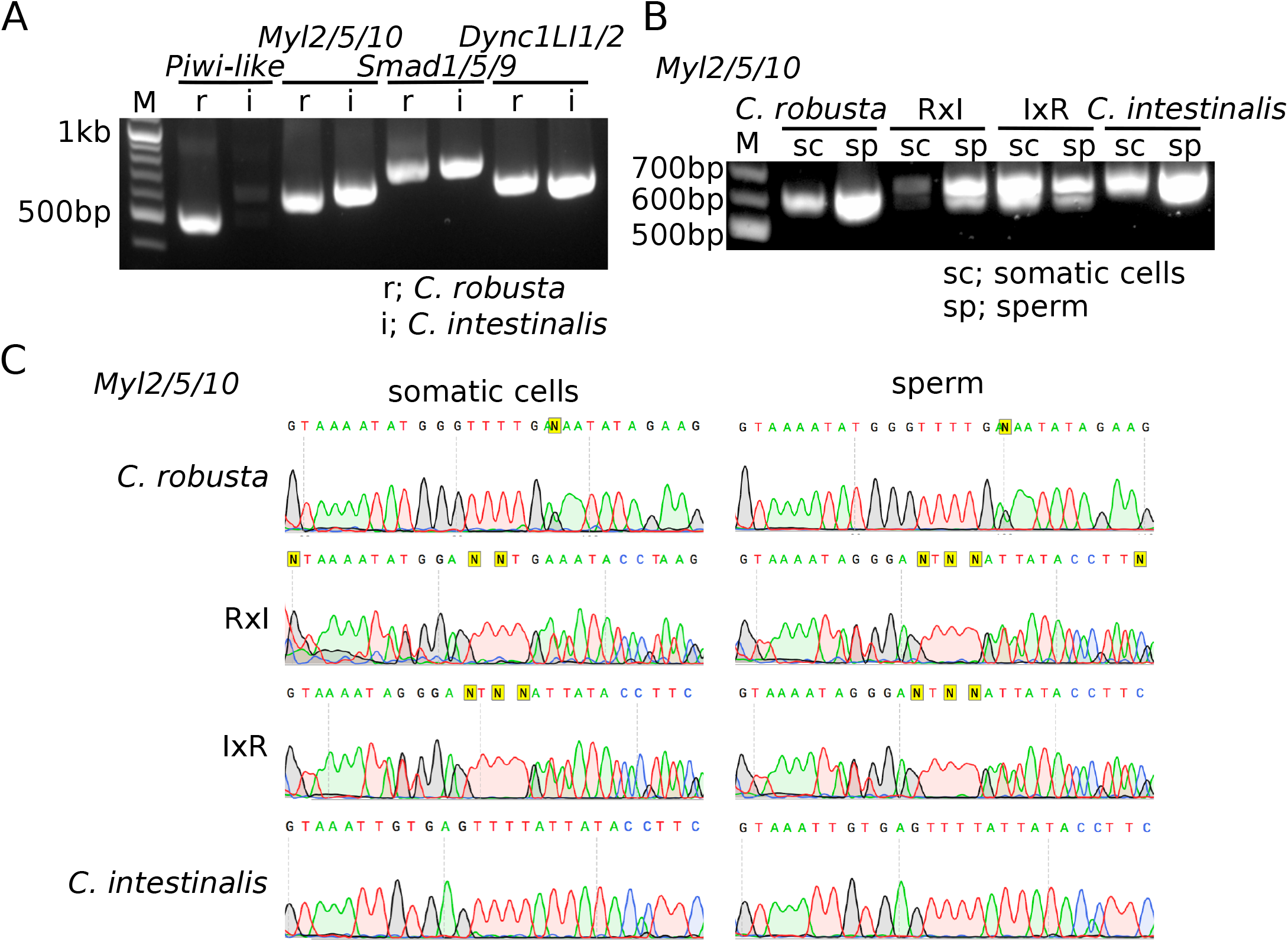
Genotype of F1 animals. (A) PCR was done with primers designed at *Piwi-like*, *Myl2/5/10*, *Smad1/5/9* and *Dync1LI1/2* gene loci from genomic DNAs from sperm of F0 mature animals of *C. robusta* and *C. intestinalis*. (B) PCR was done with primers designed at *Myl2/5/10* gene locus. Genomic DNA was collected from somatic tissue in oral siphon and sperm of F1 mature adults; *C. robusta*, RxI hybrid, IxR hybrid and *intestinalis*. (C) The sequence of PCR products in (B) were read by *Myl2/5/10*-sequence primer.

### Backcrossing to *Ciona robusta* eggs

Since we could grow F1 hybrids to sexual maturity, we sought to test whether their sperm, which appeared first, could fertilize wildtype *C. robusta* eggs. For this backcross experiment, we collected sperm from 6, 6, 8 and 6 mature F1 animals of *C. robusta*, RxI and IxR hybrids, and *C. intestinalis*, respectively (Figure 6A, Tables 1-2). On the other hand, we obtained wildtype eggs from 21 (R7-27) mature *C. robusta* animals. We crossed these sperm and eggs in 78 different combinations (summarized in Table 2). Because F2 (IxI)xR hybrids were potentially equivalent to F1 IxR hybrids, we did not analyze them further. We raised F2 *C. robusta* animals by crossing sperm from F1 *C. robusta* (RxR) animals and eggs from *C. robusta* collected from the wild, and kept F2 *C. robusta* animals as controls. We counted the proportion of fertilized eggs out of total eggs to score fertilization rates (Figure 6B-C). The fertilization rates for *C. robusta* were almost 100%, while the rates dropped and varied between 0% to 100% in the other crosses (Figure 6C). Notably, the sperm of F1 RxI hybrid appeared less potent to fertilize *C. robusta* eggs than that of F1 IxR hybrids, which is reminiscent of previously reported difficulties in using *C. robusta* eggs in interspecific fertilizations. A heatmap of the fertilization rates showed that there were no infertile eggs from wildtype *C. robusta,* while sperm from (R1I2)1, (I1R2)4 and (I1I2)2 might have been sterile (Figure 6C). Additionally, sperm which had variable fertilization rates among batches of eggs suggested sperm and egg combination-specific adverse effects on fertilization. These observations indicated that F1 hybrids of *C. robusta* and *C. intestinalis* can produce fertile sperm capable of fertilizing *C. robusta* eggs with variable efficacy, which likely constitutes a first, prezygotic, obstacle to interspecific reproduction and gene flow.

**Table 2.**
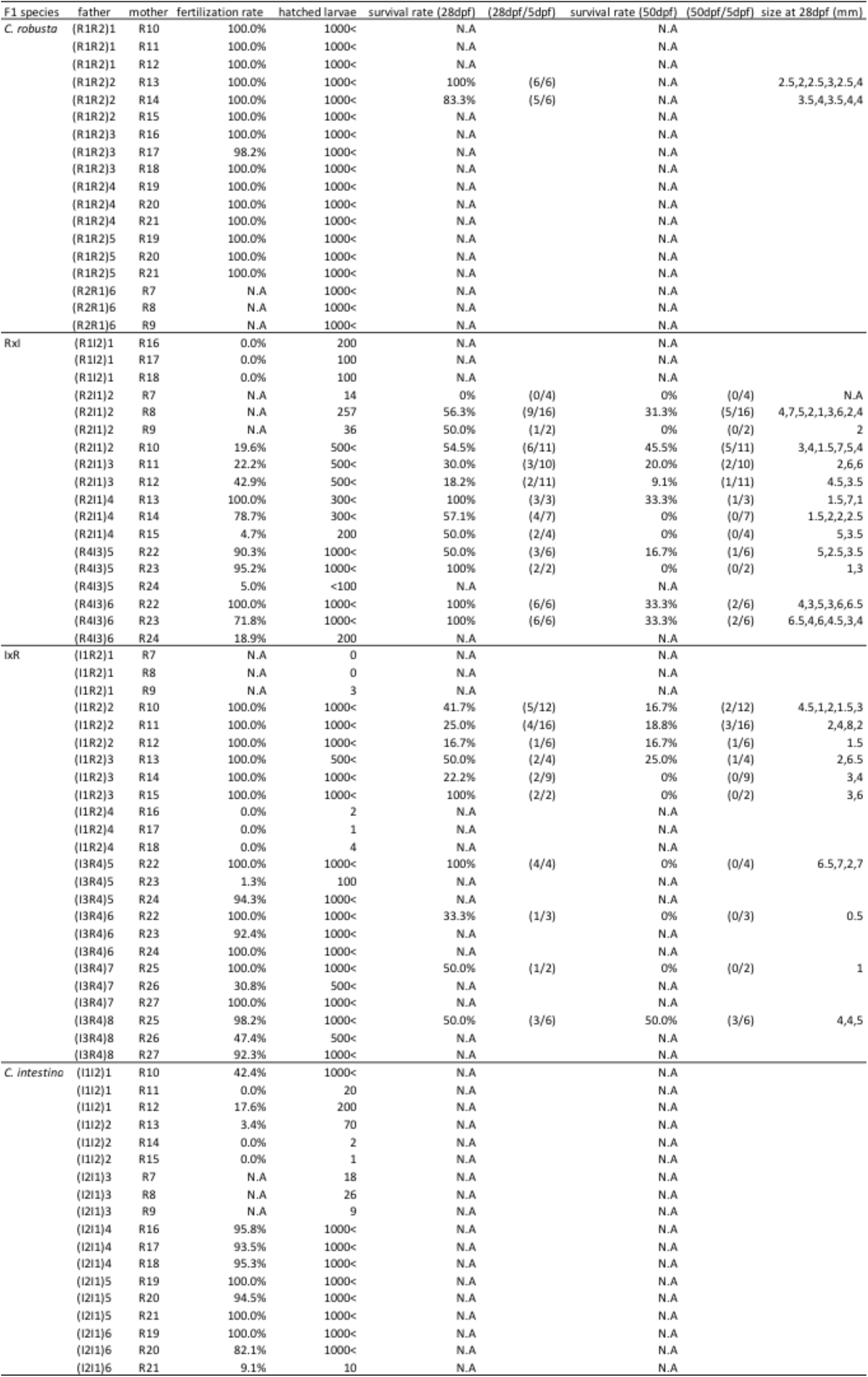
Summary BC1.

**Figure 6.**
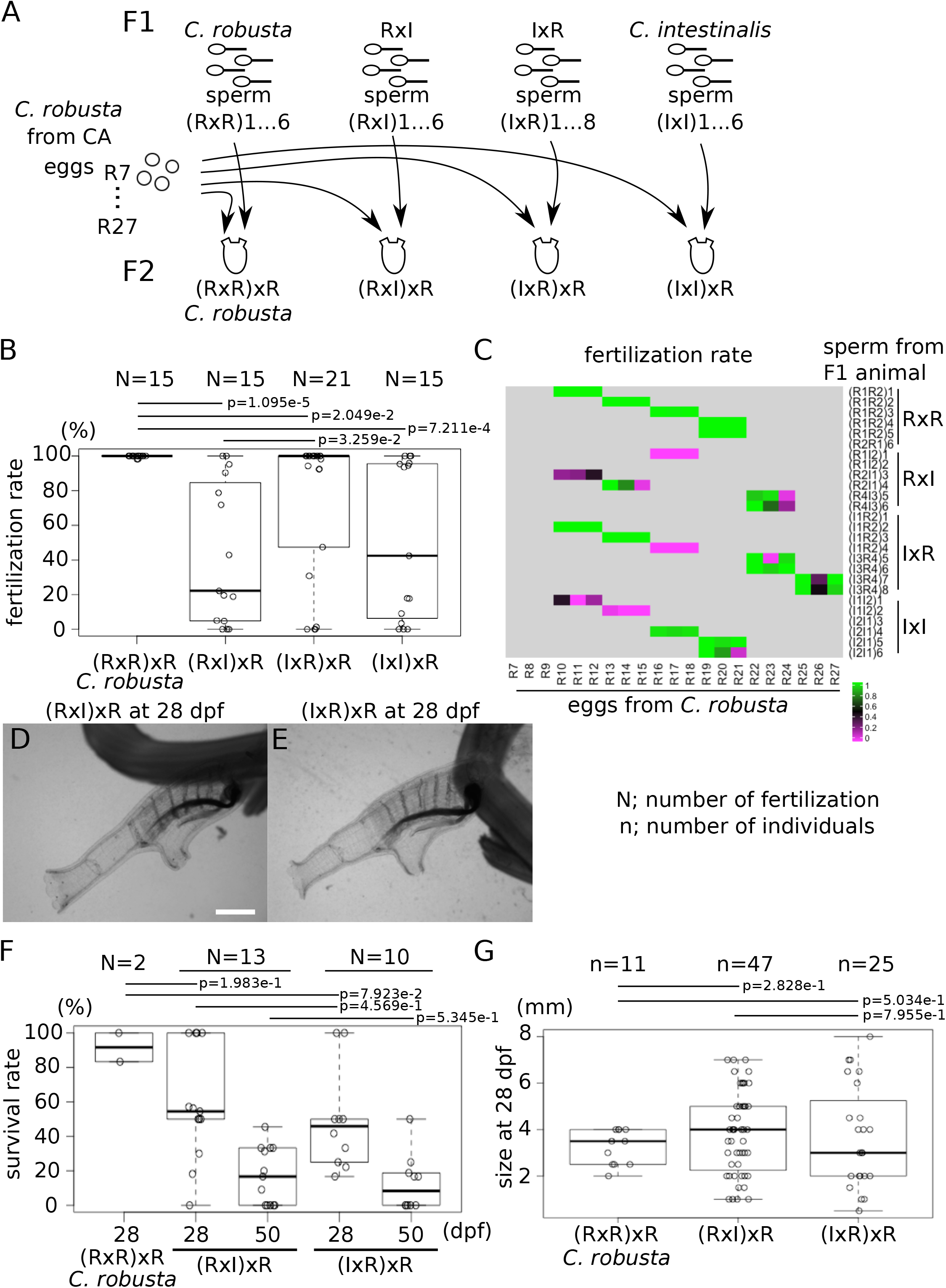
Backcrossing to *C. robusta* eggs. (A) Sperm of (RxR)1 to 6, (RxI)1 to 6, (IxR)1 to 8 and (IxI)1 to 6 were collected from F1 *C. robusta*, RxI hybrid, IxR hybrid and *C. intestinalis* mature animals, respectively. Wildtype eggs R7 to R27 were collected from mature animals in CA. (B) The dot and boxplot shows the fertilization rate. (C) The heatmap shows the fertilization rate in each fertilization. (D, E) Young adults of F2 (RxI)xR (D) and (IxR)xR (E) at 28 dpf were imaged under a microscope. Scale bar shows 1 mm. (F) The dot and boxplot shows the survival rate of each fertilization. (G) The dot and boxplot shows the size of each individual at 28 dpf. N shows the numbers of fertilization. n shows the numbers of individuals. p values were calculated by an ANOVA.

Sato and colleagues also successfully obtained mature F1 hybrids, which could be backcrossed to parental species, and the backcrossed BC1 hybrids could develop into seemingly normal larvae (Sato *et al.* 2014). Likewise, we raised BC1 (RxI)xR and (IxR)xR hybrids at 18℃, and allowed them to metamorphose and become young adults by 28 dpf (Figure 6D-E). As a measure of hybrid fitness, we calculated survival rates by counting the number of animals that survived to 28 and 50 dpf relative to the numbers of juveniles at 5 dpf (Figure 6F). Only half of BC1 (RxI)xR and (IxR)xR hybrid juveniles survived to 28 dpf, compared to almost 90% for *C. robusta*. Approximately 20% of juveniles of both BC1 hybrids survived to 50 dpf. Both BC1 (RxI)xR and (IxR)xR hybrids had lower survival rates than F2 *C. robusta* animals, while an ANOVA did not show significant differences in survival rate on 28 and 50 dpf between (RxI)xR and (IxR)xR hybrids. These observations suggest that BC1 hybrid juveniles experience higher mortality rates, consistent with proposed genomic incompatibilities in second generation hybrids (Dobzhansky-Müller Incompatibilities, DMI, (Malfant *et al.* 2018)).

As a complement to survival, we measured the body size of BC1 animals at 28 dpf (Figure 6G). The size of F2 *C. robusta* juveniles varied between 2 and 4 mm (average=3.23, SD=0.75, n=11), while the size of BC1 (RxI)xR and (IxR)xR hybrids varied between 0.5 and 8 mm ((RxI)xR; average=3.82, SD=1.78, n=47, (IxR)xR; average=3.70, SD=2.22, n=25). This suggested that growth rates are more variable in the BC1 hybrid population, as expected following the segregation of alleles for a likely multigenic trait such as individual growth rate.

Seventeen and ten individuals of (RxI)xR and (IxR)xR hybrids grew to mature adults, respectively, thus allowing us to observe the presence of OPO and the color of their siphons (Figure 7 and Table 3). Except for one individual [(R2I1)2xR8], BC1 (RxI)xR and (IxR)xR hybrids had OPO at the tip of the sperm duct (Figure 7A and Table 3), which is also consistent with the presence of OPO being a dominant *C. robusta* trait. Half of the individuals in both BC1 (RxI)xR and (IxR)xR hybrids had red color in the rim of siphons, as did F1 hybrids, while the other half had transparent siphons, the same as normal *C. robusta* (Figure 7B). This could be explained considering a single gene, with distinct *C. robusta* and *C. intestinalis* alleles, which coexist in F1 hybrids and segregate with a 1:1 ratio in the BC1 hybrid population, because animals heterozygous for *C. robusta* and *C. intestinalis* alleles should produce red-colored siphons, as seen in F1 hybrids, while homozygous *C. robusta* alleles should produce colorless siphons.

**Table 3.**
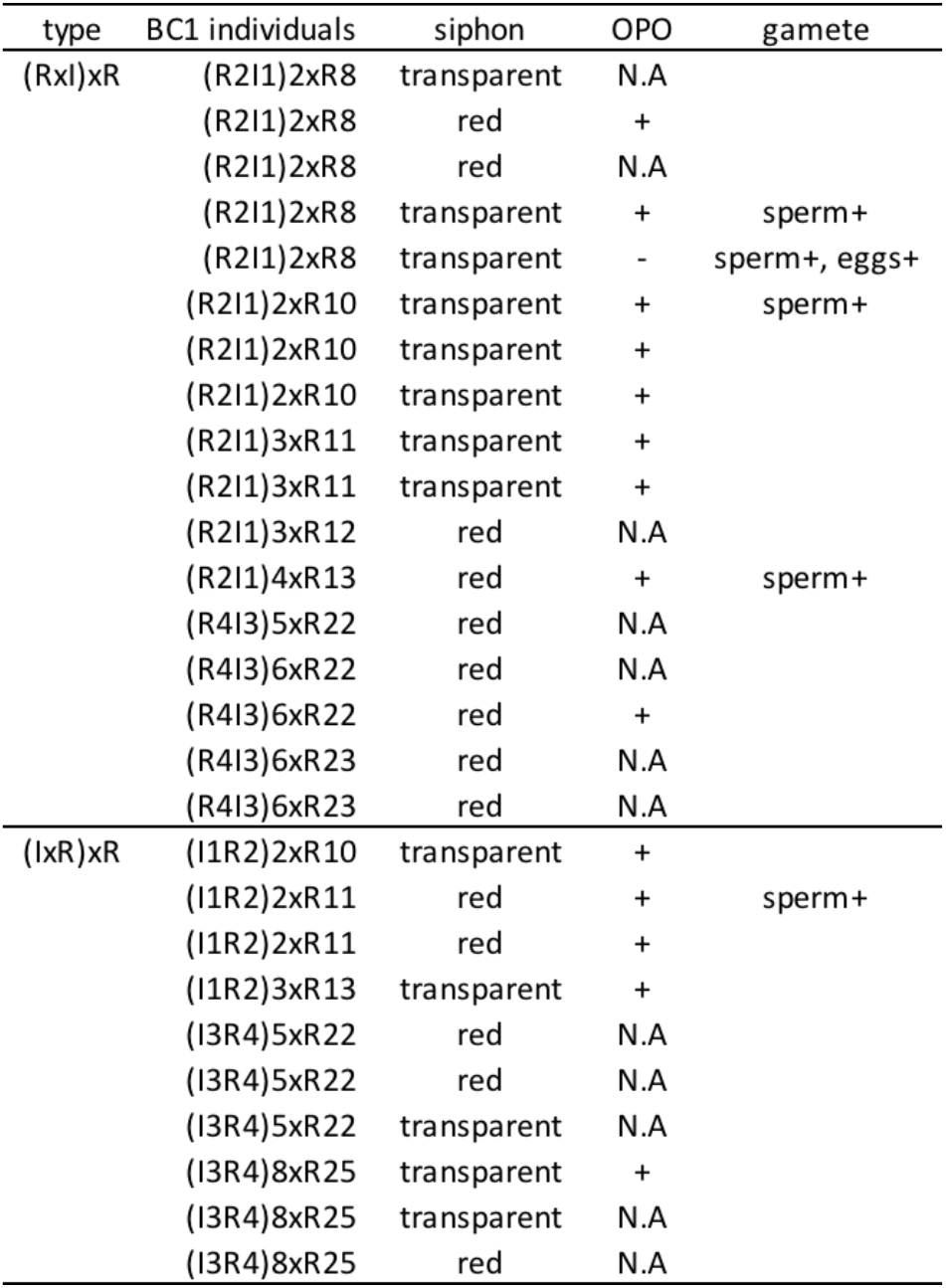
Phenotype BC1.

**Figure 7.**
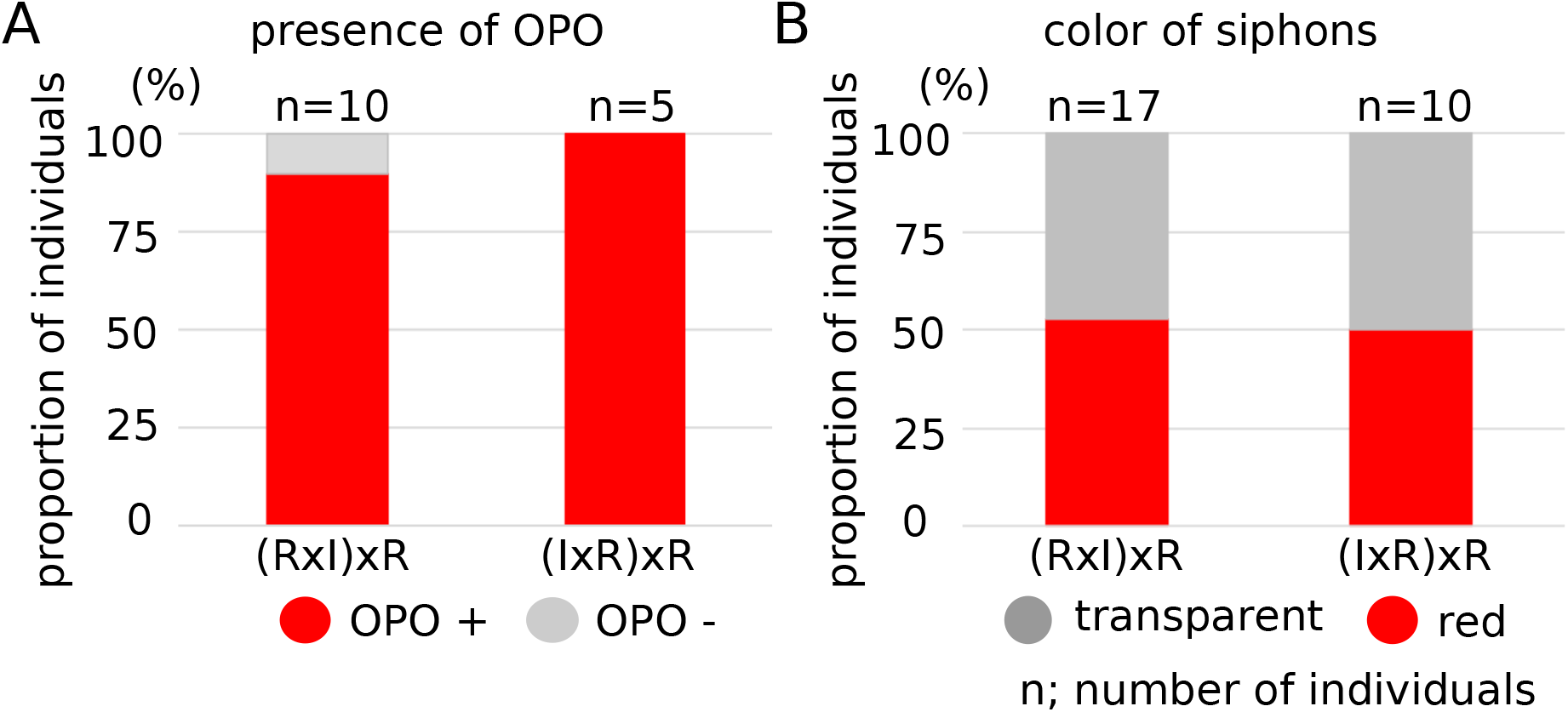
Phenotype of BC1 hybrid mature animals. (A) The bar shows the proportion of individuals having OPO at the tip of sperm duct in BC1 hybrid mature animals. (B) The bar shows the proportion of individuals having color in the rim of oral and atrial siphons in BC1 hybrid mature animals. n shows the numbers of animals.

Finally, both (RxI)xR and (IxR)xR BC1 hybrids grew and matured to produce sperm and eggs (Table 3 and Supplemental Table S2). The sperm could fertilize *C. robusta* eggs to produce BC2 hybrids, which survived at least 28 dpf, after which we stopped observations. This indicates that the BC1 hybrids that survive, grow and mature are potentially fertile. This possibility is not incompatible with the existence of the DMI. Instead, it is consistent with the existence of defined hotspots of unidirectional introgression observed in wild populations (Roux *et al.* 2013).

### Inbreeding F1 RxI and IxR hybrids

Next, we leveraged the fertility of *C. robusta* x *C. intestinalis* offspring to test whether crossing F1 hybrids would yield viable F2 animals, which would in principle provide opportunities for quantitative genetics approaches for the analysis of complex traits. We obtained sperm from 7 and 10 individuals, and eggs from 7 and 11 F1 RxI and IxR mature animals, respectively, and used them for within-type fertilizations (Figure 8A, Tables 1 and 4). Fertilization rates were significantly higher for IxR hybrids than for RxI hybrids, suggesting asymmetric second generation incompatibilities between *C. robusta* and *C. intestinalis* genotypes (Figure 8B). Specifically, crosses between IxR hybrids yielded almost 100% fertilization in 11 trials, except for two combinations, (I6R5)16x(I5R6)18 and (I4R3)17x(I6R5)16, while crosses between RxI hybrids almost invariably failed, except for the (R2I1)7x(R2I1)14 combination (Figure 8C). The data suggested that the (I6R5)16 F1 adult produced unhealthy gametes, because neither sperm nor eggs yielded productive fertilization. By contrast with backcrossing fertilizations, a limited number of eggs from F1 hybrids produced only hundreds of hatched larvae, thus limiting the numbers of F2 hybrid juveniles in each Petri dish (Table 4). Thus, we calculated metamorphosis rates of F2 hybrids by counting the number of juveniles relative to the number of swimming larvae for each fertilization, and could thus evaluate 4 and 10 fertilizations for RxI and IxR crosses, respectively (Figure 8D). The metamorphosis rates of F2 RxI and IxR hybrids ranged from 0 to 6% and 14%, respectively, which were lower than for *C. robusta* in regular fertilization (2-80%, average=25.6%, SD=15.6, N=33). Notably, an ANOVA showed significant differences in the metamorphosis rate between F2 RxI and IxR hybrids. Because of low fertilization and metamorphosis rates, we obtained only 7 F2 RxI hybrid juveniles by 5 dpf, compared to 137 F2 IxR hybrid juveniles (Table 4). In total, 3 and 85 juveniles of F2 RxI and IxR hybrids survived to 28 dpf, and displayed normal morphologies, similar to *C. robusta* (Figure 8E-F). There were no obvious morphological differences among 28 dpf F2 hybrid individuals between ross types. Survival rates were calculated by counting the number of individuals that survived to 28 dpf and 50 dpf, relative to the number of juveniles at 5 dpf (Figure 8G). Only 1 juvenile from the (R2I1)9x(R4I3)15 and (R4I3)10x(R5I6)16 crosses survived to 50 dpf, and there were only 2 individuals of F2 RxI hybrids that survived to 50 dpf, compared to 57 F2 IxR hybrid individuals, suggesting that F2 hybrids were less viable in the RxI type than in the IxR type. This intriguing observation suggested the existence of asymmetric second generation genomic incompatibilities, which could involve maternal determinants such as the mitochondrial genome. For example, it is conceivable that homozygous *C. robusta* alleles for certain loci would be incompatible with *C. intestinalis* mitochondrial DNA, as this combination would preferentially emerge in second generation RxI crosses, assuming quasi-exclusive maternal inheritance of the mitochondrial DNA (Nishikata *et al.* 1987).

**Table 4.**
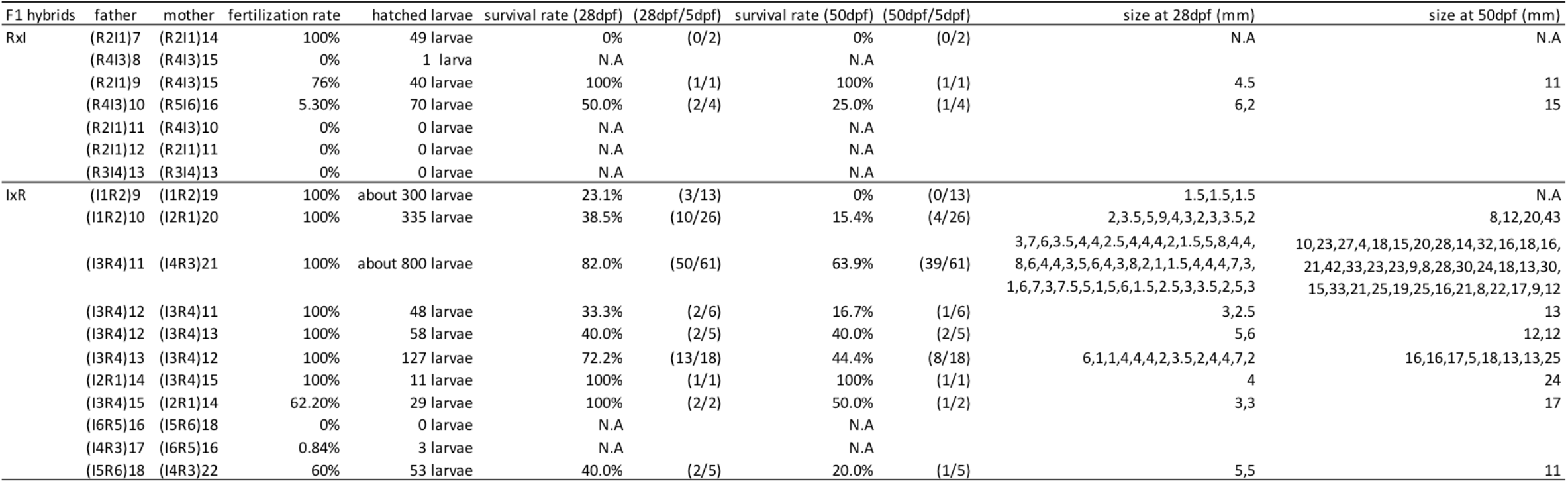
Summary inbreeding F2.

**Figure 8.**
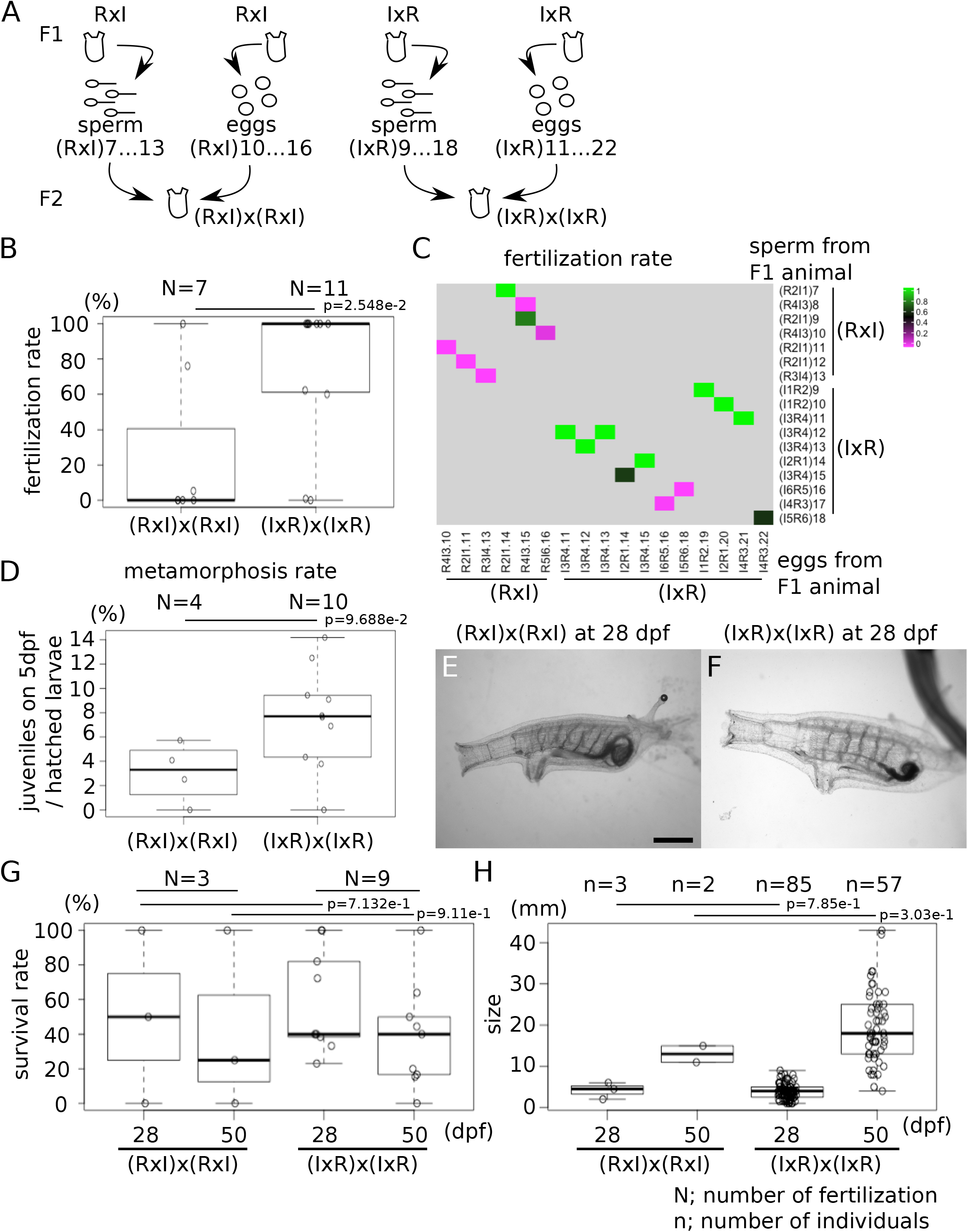
Inbreeding of F1 RxI and IxR hybrids. (A) Sperm and eggs were collected from F1 RxI hybrid of (RxI)7 to 13 and (RxI)10 to 16, respectively. Sperm and eggs were collected from F1 IxR hybrids of (IxR)9 to 18 and (IxR)11 to 22, respectively. These sperm and eggs were crossed to produce F2 hybrids. (B) The dot and boxplot shows the fertilization rate. (C) The heatmap shows the fertilization rate in each fertilization. (D) The dot and boxplot shows the metamorphosis rates. (E, F) Young adults of F2 RxI (E) and IxR (F) hybrids at 28 dpf were imaged under a microscope. Scale bar=1 mm. (G) The dot and boxplot shows the survival rate of each fertilization. (H) The dot and boxplot shows the size of individuals at 28 and 50 dpf. N shows the numbers of fertilization. n shows the number of individuals. p values were calculated by an ANOVA.

We could measure body sizes for only 3 and 2 F2 RxI hybrid individuals at 28 and 50 dpf, preventing robust statistical analysis. By contrast, 85 and 57 F2 IxR hybrid individuals measured at 28 and 50 dpf showed a range of body sizes similar to that of BC1 hybrids (Figure 8H). This is also consistent with the notion that body size is a polygenic trait, which displays increased continuous phenotypic variation following alleles segregation of multiple genes in F2. This also suggests that these animals are not obviously subjected to second generation genomic incompatibilities, at least in IxR crosses. Finally, although body size is likely multifactorial, i.e. influenced by the environment, especially the availability of food, we surmise that most of the observed variation in controlled laboratory conditions is due to polygenic effects.

Following Mendel’s laws, the proportions of homo- and heterozygous animals among F2 hybrids should follow a 1:2:1 distribution in the absence of hybrid dysgenesis, inbreeding depression and/or second generation genomic incompatibilities. We analyzed the genotypes at *Myl2/5/10* and marker 2 loci for 24 swimming larvae in each of two lines of F2 IxR hybrids, ((I1R2)10x(I2R1)20 and (I3R4)11x(I4R3)21) (Figure 9). At the *Myl2/5/10* locus, there were 4 and 6 larvae showing homozygous *C. robusta* alleles, 12 and 13 heterozygous larvae, and 1 and 2 larvae homozygous for the *C. intestinalis* allele out of 17 and 21 verified samples in (I1R2)10x(I2R1)20 and (I3R4)11x(I4R3)21, respectively (Figure 9E). The proportion of *C. intestinalis* genotype was significantly different from the theoretical estimation 25% (p=1.354e-2 by z-test). By contrast, at the marker 2 locus, there were 2 and 1 larvae homozygous for the *C. robusta* allele, 13 and 18 heterozygous larvae, and 9 and 5 larvae homozygous for the *C. intestinalis* allele out of 24 and 24 verified samples in (I1R2)10x(I2R1)20 and (I3R4)11x(I4R3)21, respectively (Figure 9F). At this locus, the proportion of *C. robusta* genotype was significantly different from the theoretical estimation 25% (p=1.286e-3 by z-test). Biased genotype in *Myl2/5/10* showing less *C. intestinalis* type and marker 2 showing less *C. robusta type*, suggests that these genes of homozygous type are linked to loci causing genomic incompatibility in F2 hybrids populations. Because *Myl2/5/10* and marker 2 genes are on different chromosomes and neither are located in the inferred hotspots of introgression (Roux *et al.* 2013), their allelic distributions might be independent and differentially affected by linkage with incompatible loci. Future work will be required to characterize the genetic underpinnings of genomic incompatibilities between *Ciona* species, their relationships to documented “hotspots” of introgression (Roux *et al.* 2013), and their impact on speciation.

**Figure 9.**
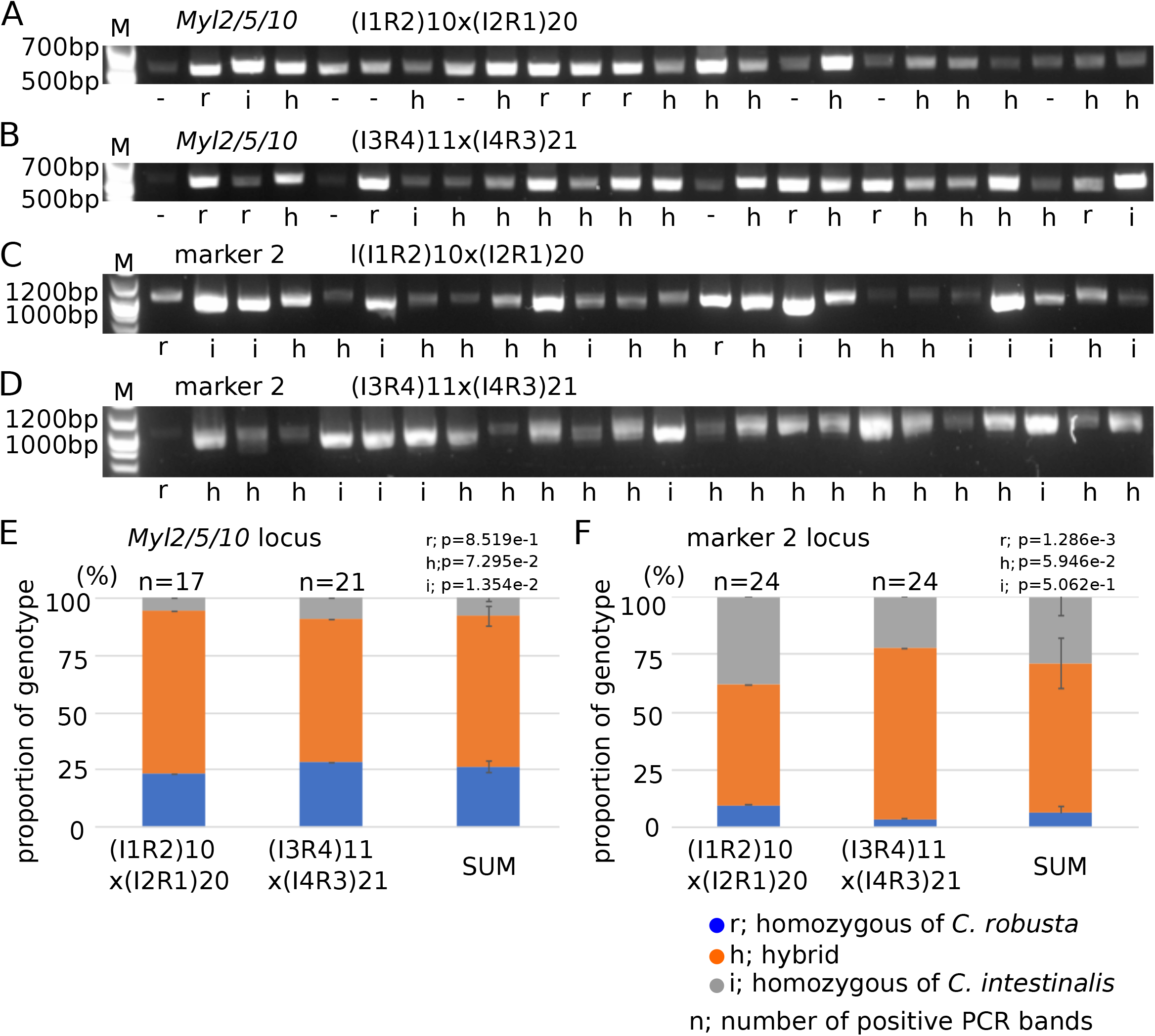
Genotyping of F2 IxR hybrid. (A-D) PCR was done with primers designed at *Myl2/5/10* (A, B) and marker 2 (Suzuki *et al.* 2005) gene loci. Genomic DNAs were collected from F2 larvae; each 24 larva from (I1R2)10x(I2R1)20 (A, C) and (I3R4)11x(I4R3)21 (B, D). (E, F) The bar shows the proportion of genotypes at *Myl2/5/10* (E) and marker 2 (F) gene loci in (I1R2)10x(I2R1)20 and (I3R4)11x(I4R3)21, and sum of two. p values were calculated by z-test from theoretical predicted value, 25% or 50%.

## Discussion

In this study, we crossed the ascidian species *Ciona robusta* and *Ciona intestinalis* to establish hybrid lines, further probe the reproductive isolation of these recently distinguished species, and explore opportunities for quantitative genetics using the genome-enabled *Ciona* model. Taking advantage of a simple closed inland culture system, we monitored post-embryonic development and survival, and successfully raised F1 and F2 hybrid and backcrossed animals to maturity. The partial viability of first and second generation hybrids provided insights into the genetics of simple traits, such as the presence of OPO, which appears to be a dominant *C. robusta*-specific trait. On the other hand, siphon pigmentation showed an overdominant phenotype in hybrids, suggesting more complex genetic interactions, although the distribution in F2 hybrids could be explained by allele segregation at one locus. Moreover, simple quantitative traits, such as body size, showed an increased variability in F2 hybrids as expected for polygenic traits following allele segregation. These observations suggest that quantitative genetics approaches could be used to study complex traits that differ between *C. intestinalis* and *C. robusta*, such as tolerance to high water temperature (Caputi *et al.* 2015; Malfant *et al.* 2017).

Despite preliminary evidence of allele segregation in F2 hybrids, the representation of genotype combinations is likely to be biased due to genomic incompatibilities, which might hinder quantitative analysis of polygenic traits. Indeed, our observations suggest that both pre- and postzygotic mechanisms contribute to genomic incompatibilities in hybrids, and thus act as obstacles to interspecific reproduction between these two *Ciona* species. These observations are consistent with previous reports (Nydam and Harrison 2011; Sato *et al.* 2012; Bouchemousse *et al.* 2016a; c). Nonetheless, the incomplete penetrance of first and second generation incompatibilities suggests that certain combinations of *C. robusta* and *C. intestinalis* genotypes are viable, which would permit at least low levels of gene flow between populations, and is consistent with the existence of previously reported hotspots of introgression (Roux *et al.* 2013).

Asymmetric fertilization success in reciprocal interspecific crosses (Turelli and Moyle 2007) was previously observed between *Ciona robusta* and *Ciona intestinalis* (Suzuki *et al.* 2005; Bouchemousse *et al.* 2016a; Malfant *et al.* 2018), but the mechanisms remain elusive. On the other hand, asymmetric second generation inviability and infertility between hybrids suggested that mechanisms of genomic incompatibility involve interactions between the zygotic and/or maternal genomes and maternally inherited determinants, such as mitochondrial DNA (Turelli and Moyle 2007; Burton and Barreto 2012; Sloan *et al.* 2017). For instance, incompatibilities between the nuclear and mitochondrial genomes in hybrids were reported in various organisms, including fungi (Lee *et al.* 2008; Presgraves 2010), insect (Meiklejohn *et al.* 2013; Hoekstra *et al.* 2013), nematode (Chang *et al.* 2016) and mammals (Ma *et al.* 2016). Finally, it is tempting to speculate that these asymmetric incompatibilities provide a mechanistic explanation for the unidirectional introgressions observed in wild populations (Roux *et al.* 2013).

## Acknowledgments

We are indebted to David Remsen, Josh Rosenthal, Dana Mock-Muñoz de Luna and staff members at MBL, Woods Hole, for extensive support. We thank Régis Lasbleiz for installing the original culture system in our laboratory, and providing detailed instructions for algae and animal culture. We are grateful to Matthew Rockman for critical reading of the manuscript and precious suggestions, insightful discussions and help with the evolutionary theory, concepts and methods of quantitative and population genetics. This research was supported by NIH/NIGMS award R01 GM096032 and by an MBL Whitman Fellowship to L.C.

---

**Supplemental Figure S1.** A closed inland culture system. (A-C) Animals were cultured in 20L tank (A), polypropylene 5L beaker (B) and 2L aquarium tank (C). (D) The bar shows the survival rate of animals. (E) The bar shows the proportion of animals having sperm. n shows the numbers of individuals.

**Supplemental Figure S2.** Genotyping of F1 juveniles at marker 2 gene locus. (A) PCR was done with primers designed at marker 1 and marker 2 gene loci (Suzuki *et al.* 2005) from genomic DNAs of F0 *C. robusta* and *C. intestinalis* sperm. (B) Three F1 juveniles in each type were analyzed by PCR with marker 2 primers. (C) PCR was done with marker 2 primers with genomic DNAs collected from oral siphon and sperm of F1 *C. robusta*, RxI hybrid, IxR hybrid and *intestinalis* mature animals. (D) Sequences were read from the PCR products.

**Supplemental Figure S3.** Genotyping of F1 juveniles at *Myl2/5/10* gene locus. (A, B) PCR was done from genomic DNA of F0 *C. robusta* (A) and *C. intestinalis* (B) sperm. (C) Alignment of *Myl2/5/10* sequence between *C. robusta* and *intestinalis*. Cian shows exon sequence. Arrow shows sequence primer. Underlines show the region of sequence in Figures 5C, S2D and S3E. (D) Three F1 juveniles in each strain were analyzed by PCR with *Myl2/5/10* primers. (E) Sequences were read from the PCR products.

