## Supplemental Materials for "Asymmetric second-generation genomic incompatibility in interspecific crosses between *Ciona robusta* and *Ciona intestinalis*"

### Supplemental Figure.1 Inland culture System

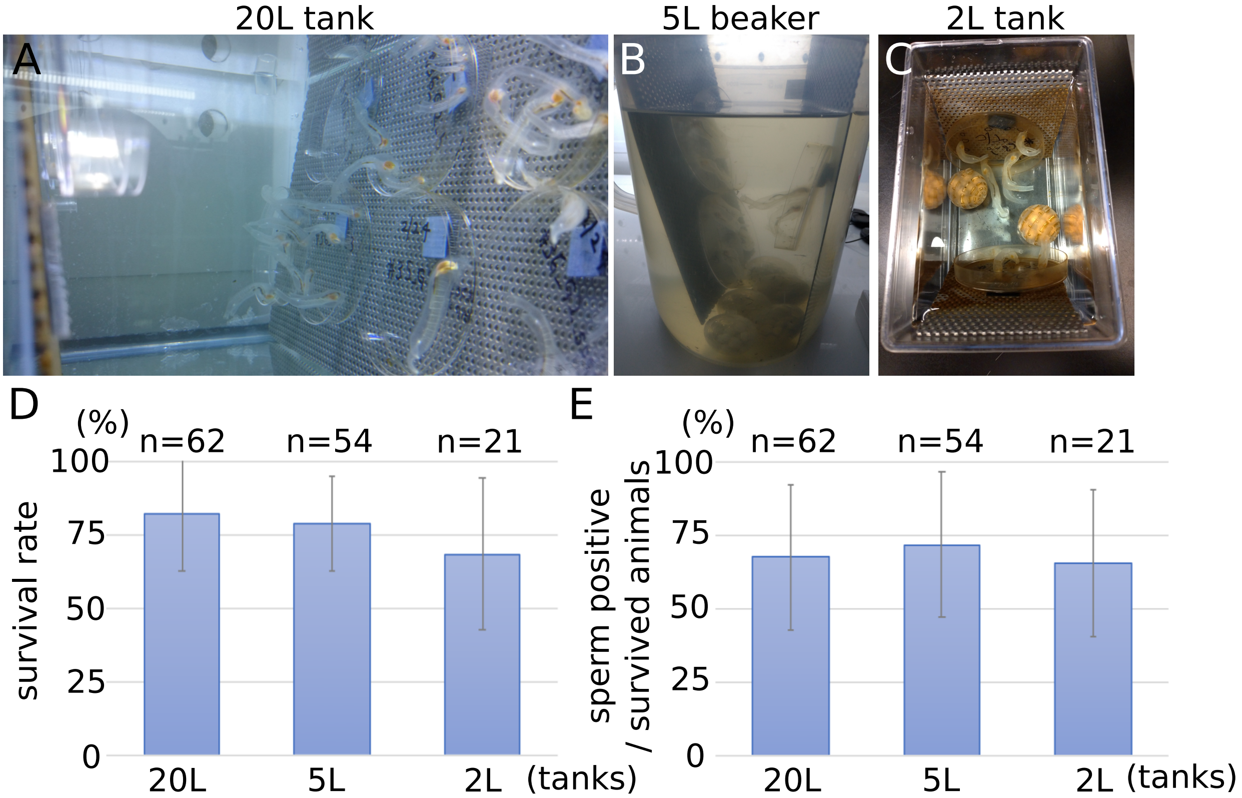


n; number of individuals

### Supplemental Figure.2 M2 locus

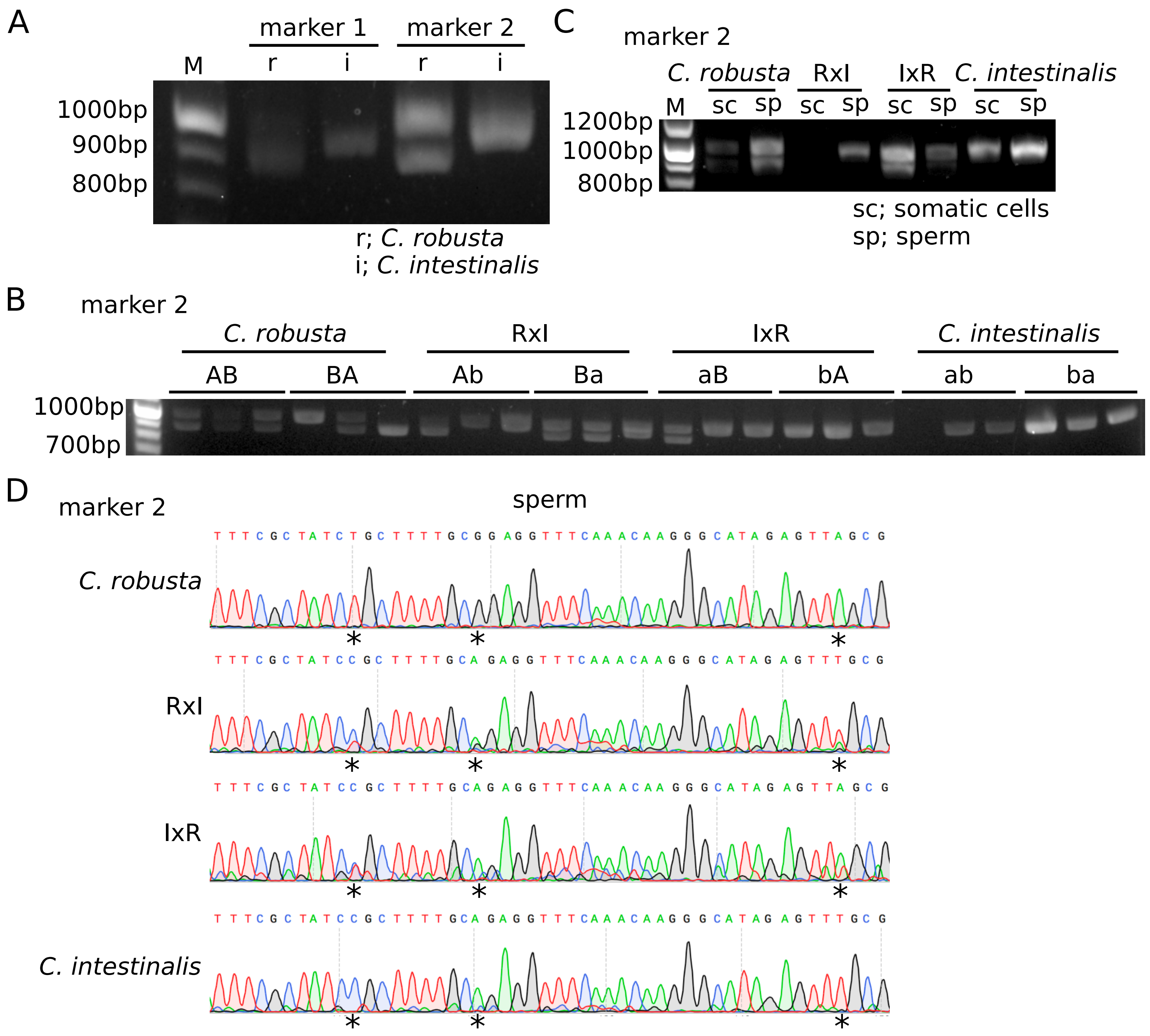

Supplemental Figure.3 F1 genotyping MYL

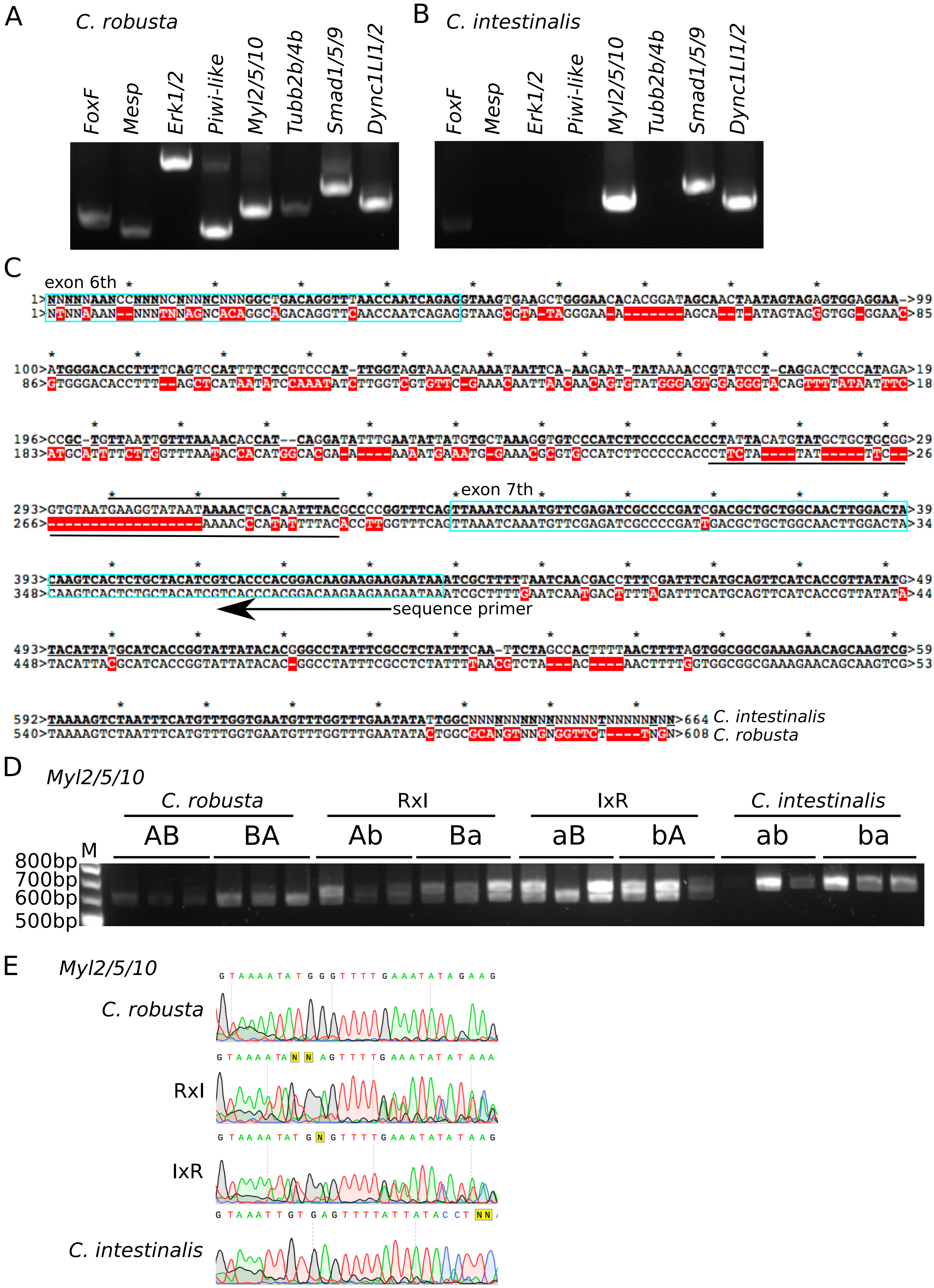

### Supplemental Table.S1 Primers

| gene name | | primers sequence |
|---|---|---|
| *FoxF* | Forward | 5'-GAATTCCGTCCGCCACAATC-3' |
| | Reverse | 5'-AGTGGAAACCTCAGCACAGT-3' |
| *Mesp* | Forward | 5'-TGTTTTGGGGATAAATGGGGA-3' |
| | Reverse | 5'-GACACACATTGGTTAACGGGT-3' |
| *Erk1/2* | Forward | 5'-CCGTTACTACACTTGTTTTCGC-3' |
| | Reverse | 5'-ACTCATGGGCGCTTTGATTT-3' |
| *Piwi-like* | Forward | 5'-GGAACCATTCGTGTACCAGC-3' |
| | Reverse | 5'-GTTTCTATTGCCACCCACGG-3' |
| *Myl2/5/10* | Forward | 5'-GAAATCCTCACGACACAGGC-3' |
| | Reverse | 5'-CAAGAACACAAAACCTGCGC-3' |
| | Sequence | 5'-TTCTTCTTGTCCGTGGGTGA-3' |
| *Tubb2b/4b* | Forward | 5'-TTGGCGAGATTTTCAGGTGC-3' |
| | Reverse | 5'-CCTGTGTGCATATTCCTTGGT-3' |
| *Smad1/5/9* | Forward | 5'-GTCGATCATTTTGGCCAGCT-3' |
| | Reverse | 5'-GGATCGGCGTGTCAAAATCA-3' |
| *Dync1LI1/2* | Forward | 5'-CACCAGGAAACATGTGCAAGA-3' |
| | Reverse | 5'-GCGATGGACTGAAACACACA-3' |
| Marker 1 | Forward | 5'-TGTATTTACTATTTTCAACG-3' |
| (Suzuki et al., 2005, J. Mol. Evol.) | Reverse | 5'-CTCCACTGCTAGCAACTGC-3' |
| Marker 2 | Forward | 5'-GTGTCTTAGAAGAGGAGAAT-3' |
| (Suzuki et al., 2005, J. Mol. Evol.) | Reverse | 5'-CTTTGTCCACTGAGAGCACT-3' |

### Supplemental Table.S2 Sumary BC2

| BC1 type | father | mother | fertilization rate | hatched larvae | survival rate (28dpf) | (28dpf/5dpf) | size at 28dpf (mm) |
|---|---|---|---|---|---|---|---|
| (RxI)xR | (R2I1)2xR10 | R28 | 25% | 1000< | 66.7% | (2/3) | 1.5,7 |
| | | R29 | 100% | 1000< | 66.7% | (4/6) | 1.5,3,4,5 |
| | | R30 | 94.90% | 1000< | 100% | (3/3) | 2.5,4,5 |
| (IxR)xR | (I1R2)2xR11 | R28 | 100% | 1000< | 91.7% | (11/12) | 2.5,5,5,5.5,6,7,2.5,3.5,6,1.5,3 |
| | | R29 | 100% | 1000< | 83.3% | (10/12) | 2,5.5,5,5,7,5,4.5,2,5,6.5 |
| | | R30 | 97.30% | 500< | 83.3% | (5/6) | 2,1.5,7,4,1.5 |
